# Physiological factors of importance for load carriage

**DOI:** 10.1101/157891

**Authors:** Manne Godhe, Torbjörn Helge, C. Mikael Mattsson, Örjan Ekblom, Björn Ekblom

## Abstract

The energy expenditure during carrying no load, 20, 35 and 50 kg at two walking speeds, 3 and 5 km h^-1^, was studied in 36 healthy participants, 19 men (30 ± 6 yrs, 82.5 ± 7.0 kg) and 17 women (29 ± 6 yrs, 66.1 ± 8.9 kg). Anthropometric data, leg muscle strength as well as trunk muscle endurance and muscle fibre distribution of the thigh were also obtained. To load the participant a standard backpack filled with extra weight according to the carrying weight tested was used. Extra Load Index (ELI), the oxygen uptake (VO_2_) during total load over no-load-exercise, was used as a proxy for load carrying ability. In addition to analyzing factors of importance for the ELI values, we also conducted mediator analyzes using sex and long term carrying experience as causal variables for ELI as the outcome value.

For the lowest load (20 kg), ELI20, was correlated with body mass but no other factors. Walking at 5 km h^-1^ body mass, body height, leg muscle strength and absolute VO_2_max were correlated to ELI35 and ELI50, but relative VO_2_max, trunk muscle endurance and leg muscle fibre distribution were not.

Sex as causal factor was evaluated in a mediator analyses with ELI50 as outcome. ELI50 at 5 km h^-1^ differed between the sexes. The limit for acceptable body load, 40 % of VO_2_max (according to Åstrand, 1967), was nearly reached for women carrying 35 kg (39%) and surpassed at 50 kg at 3 km h^-1^, and for men carrying 50 kg at 5 km h^-1^. This difference was only mediated by difference in body mass. Neither muscle fibre distribution, leg muscle strength, trunk muscle endurance and body height nor did absolute or relative VO_2_max explain the difference.

Participants with long term experience of heavy load carrying had significant lower ELI20 and ELI50 values than those with minor or non-experience, but none of the above studied factors could explain this difference.

The study showed that body mass and experience of carrying heavy loads are important factors for the ability to carry heavy loads.

**Funding:** This study was founded by the Swedish Military Forces´ Research Authority and The Research Funds of The Swedish School of Sport and Health Sciences.

**Abbreviations:**

ELI: Extra Load Index
HLa: Blood lactate concentration
HR: Heart rate
RPE: Rate of Perceived Exertion
VO_2_: Oxygen uptake
VO_2_max: Maximal oxygen uptake

## Introduction

To be able to carry heavy loads is an important and necessary task during many outdoor physical activities, especially during military operations. Numerous studies have analyzed the physiological and biomechanical consequences of different loads and speeds. Most of them have shown that energy cost of walking increases progressively with both load and walking speed (Goldman and Lampietro, 1962, Pandolf et al. 1976, Epstein et al. 1998, Bastien et al. 2005). Christie and Scott (2005) showed that the highest combination of load and speed for acceptable load on the individual, evaluated as taxing the oxygen transport system to lesser than 40 % of maximal oxygen uptake (VO_2_max) as suggested by Åstrand (1967), was 35 kg at 3,5m h^-1^ and 20 kg at 4,5 km h^-1^. However, in many real life situations the loads can be much higher than that.

With regard to differences between the sexes it is generally accepted that women in a given situation experience higher average strain on oxygen transport system and higher perceived exertion than men, mainly due to the lower body mass and lesser absolute VO_2_max (l min^-1^). However, other factors of importance for load carrying ability in both sexes, when for instance women and men have equal body weight or the same relative VO_2_max (ml min^-1^ kg^-1^), have not been evaluated. Furthermore, long term experience of carrying heavy loads is evidently of importance. Sherpas and Tibetan Highlanders can carry loads > 100 % of body mass during long walks. This remarkable carrying ability can partly be explained by the high mechanical efficiency due to a better load balancing than corresponding in control subjects (Bastien et al, 2005, Marconi et al. 2005, Minetti et al. 2006). To our knowledge several factors that might be involved in load carrying analyses have not been studied.

Therefore, we studied participants both with long term experience and those with minor experience of load carrying during walking at two normal speeds (3 and 5 km h^-1^), carrying different loads (20, 35 and 50 kg). Oxygen uptake (VO_2_), anthropometric data, leg muscle strength as well as trunk muscle endurance and muscle fibre distribution of the thigh were obtained. In addition we compared VO_2_ during load walking with VO_2_ during no load walking at the two speeds. Thus, a quote of VO_2_ during total load over no load walking, Extra Load Index (ELI), could be calculated. This quotient is accepted as a proxy for load carrying capacity (Taylor et al. 1980, Lloyd et al. 2010). Furthermore, in addition to analyze factors of importance for the carrying ability (ELI values) we also conducted mediator analyzes using sex and long term carrying experience as causal variables for ELI as the outcome value.

## Methods and procedures

### Participants

Thirty six volunteers, nineteen men (30 ± 6 years, 82.5 ± 7.0 kg) and seventeen women (29 ± 6 years, 66.1 ± 8.9 kg), provided written informed consent to participate in the study after being verbally informed. Inclusion criteria was healthy status with no current or past injury that could compromise participation. For anthropometric and physiological data – see Table 1. Half of the participants were recruited from firefighters, military personnel and Special Force Police officers, and the other half from The Swedish School of Sport and Health Sciences. Both groups included participants with long term experience (> 5 years) from carrying heavy loads (n = 16, 8 women) or had minor or no such experience (n = 20, 9 women). The study was approved by the Regional Ethical Review Board in Stockholm.

**Table 1.**
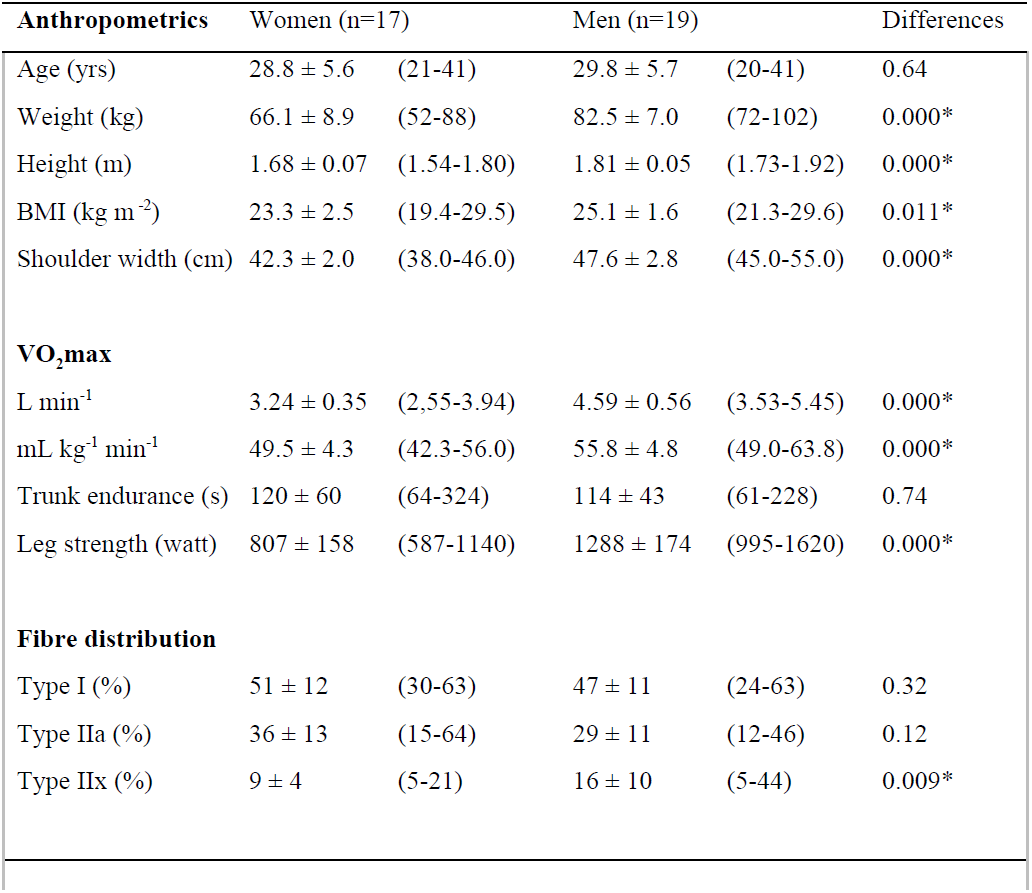
Anthropometric and other data in women and men (mean, SD, range and statistical difference between the sexes). *denotes p<0.05.

| **Anthropometrics** | Women (n=17) |  | Men (n=19) |  | Differences |
| --- | --- | --- | --- | --- | --- |
| Age (yrs) | 28.8 ± 5.6 | (21-41) | 29.8 ± 5.7 | (20-41) | 0.64 |
| Weight (kg) | 66.1 ± 8.9 | (52-88) | 82.5 ± 7.0 | (72-102) | 0.000* |
| Height (m) | 1.68 ± 0.07 | (1.54-1.80) | 1.81 ± 0.05 | (1.73-1.92) | 0.000* |
| BMI (kg m<sup>-2</sup>) | 23.3 ± 2.5 | (19.4-29.5) | 25.1 ± 1.6 | (21.3-29.6) | 0.011* |
| Shoulder width (cm) | 42.3 ± 2.0 | (38.0-46.0) | 47.6 ± 2.8 | (45.0-55.0) | 0.000* |
| **VO<sub>2</sub>max** |  |  |  |  |  |
| L min<sup>-1</sup> | 3.24 ± 0.35 | (2.55-3.94) | 4.59 ± 0.56 | (3.53-5.45) | 0.000* |
| mL kg<sup>-1</sup> min<sup>-1</sup> | 49.5 ± 4.3 | (42.3-56.0) | 55.8 ± 4.8 | (49.0-63.8) | 0.000* |
| Trunk endurance (s) | 120 ± 60 | (64-324) | 114 ± 43 | (61-228) | 0.74 |
| Leg strength (watt) | 807 ± 158 | (587-1140) | 1288 ± 174 | (995-1620) | 0.000* |
| **Fibre distribution** |  |  |  |  |  |
| Type I (%) | 51 ± 12 | (30-63) | 47 ± 11 | (24-63) | 0.32 |
| Type IIa (%) | 36 ± 13 | (15-64) | 29 ± 11 | (12-46) | 0.12 |
| Type IIx (%) | 9 ± 4 | (5-21) | 16 ± 10 | (5-44) | 0.009* |

### Design and experimental protocol

The participants performed different tests at a total of five different occasions on separate days with at least 1 day interval. The tests included a reference test, one unloaded and three loaded (20, 35 and 50 kg) walking tests at two speed (3 and 5 km h^-1^). At the fifth occasion different strength tests were carried out and in addition muscle tissue was biopsied from the lateral part of the Quadriceps muscle. For the reference test a treadmill (Rodby, Södertälje, Sweden) was used. The unloaded and loaded walk tests were carried out on carpets positioned in a rectangular shaped path (total length of 30 m per lap), consistent of 5 cm high massive soft material and included two equal blocks of 35 cm height, 135 cm length and 50 cm width, mimicking outdoor walking in varying terrain.

### Reference test

The aim of the reference test was to determine VO_2_, heart rate (HR) and blood lactate concentration (HLa) during standstill and at different submaximal rates of work. Immediately after each submaximal exercise bout a fingertip blood sample for determination Hla and RPE for general, back, leg and breathing fatigue were obtained. Finally, a conventional VO_2_max test with stepwise increasing speed and inclination was carried out during which maximal values were obtained.

### Walking tests

To load the participant a standard backpack filled with extra weight according to the carrying weight tested was used. At the start of the experiment the participants stood still for two minutes. Before one of the three occasions the participants walked at 3 and 5 km h^-1^ without carrying any extra load. At the loaded tests they walked for 5 min at 3 and 5 km h^-1^ with one minute of intermittent rest carrying either 20, 35 or 50 kg in a random order. One female participant could not finish the 50 kg, 5 km h^-1^ test due to fatigue. During the 5 min walk the participants completed 9 laps during the 3 km h^-1^ walk and 14 laps during the 5 km h ^-1^ walk. The speed was paced via lights and signals using wireless lamps (Fitlight, Fitlight sports corporation, Aurora Ontario, Canada). The measurements of VO_2_ and other parameters during carpet walking were exactly the same as during the reference test. During all tests the participants carried three lightweight (27 g) electronic accelerometers, model GT3X+ (AntiGraph LCC, Pensacola, FL, USA), one on the hip, one on the chest and one on the wrist of the dominant hand in order to measure movements in three directions (x-, y- and z-directions) and also summarize the count in a general vector count on the three places.

### Methods

Body height and weight were measured before each experiment days using standardized methods to the nearest 0.1 cm and 0.1 kg, respectively.

VO_2_ during the reference test was measured with a stationary (Oxycon Pro) and during the carpet walking with a mobile online system (Oxycon mobile), both from Erich Jaeger GmbH, Hoechberg, Germany. These systems have been validated against the Douglas bag-method (Rietjens, et al. 2001, Rosdahl et al. 2010) and also against each other without any notable differences (Akkermans et al. 2012). The fingertip blood HLa was measured using a laboratory standard method (Biosen C-line, EKF Diagnostic, Barleben Germany). HR was continuously measured with a heart rate monitor (RS800, Polar Electro Oy, Finland). The RPE was evaluated according to Borg (Borg 1970).

### Gross energy efficiency

The quotient between the VO_2_ when walking with loaded weight in reference to VO_2_ when walking unloaded was used as a proxy of carrying ability. It is an approach based on the equations of Taylor et al. (1980). The equation was later expressed in a simpler form to produce an index (ELI, Extra load index) by Lloyd et al. (2010). This index allows for comparison of relative gross energy efficiency during load carriage.

### Strength tests

The participants started with a short warm-up session on a bicycle ergometer. Thereafter leg muscle strength was measured with loaded strength in a Smith-machine (Cybex international Medway, USA). An explosive squat with loads of 20, 35, 50 kg and the same load as body weight were carried out. Before the test started the participants’ shoulder width was measured and marked on the floor. The participant was instructed to stand with feet wider than these floor marks. The participant dropped 30 cm from upright position and directly pushed upward. There were two separate tries on each weight with a few minutes of rest between. The highest result from either test was noted. All results were recorded with Muscle lab software and noted as power output in Watt.

Trunk muscle endurance was standardized according to Wyss et al. (2007), which is a simplified method of Tshopp et al. (2001). The participants was positioned horizontally on elbows and knees on the floor with 90° angle in both shoulder, elbow and hip joints with thumbs pointing up. The participants pressed their lower back against a bar construction. In this position participants were instructed to lift one alternating foot 5 cm above the floor every second. A metronome was used for 60 Hz tempo aid. When the participant no longer was able to hold the position with the lower back pressed against the bar, the test was terminated. The finish time was registered as seconds.

### Muscle biopsy

After a small incision through skin and fascia a Weil Blakesley conchotome (Wisex, Mölndahl, Sweden) was used to extract 75-100 mg of muscle tissue from Vastus Lateralis of the Quadriceps muscle under local anesthesia with Carbocain without epinephrine (AstraZeneca, Södertälje, Sweden) according to Ekblom (2016). The histochemical determination of muscle fibre type distribution was done with standardized laboratory methods.

### Statistics

All statistical tests were performed in Statistica 12 (StatSoft Inc. Tulsa, OK, USA). All variables were normally distributed, except for trunk endurance and distribution of type IIx muscle fibres. Differences between sexes were performed using independent t-tests, Mann-Whithey U-test and chi-square, for normally, non-normally distributed and categorical variables, respectively.

Collinearity was assumed if Spearman correlations between variables exceeded rho = 0.7.

Most of the candidate factors were highly correlated. Hence no linear regression could be performed. We applied mediation analysis according to the model proposed by Preacher and Heyes (2008) to investigate the possible mediators for sex difference and difference between participants with or without long term experience of load carrying. We tested candidates for mediation one by one, reporting bootstrapped paths coefficients and the corresponding bias adjusted 95% CIs.

## Results

### Anthropometrics

In Table 1 anthropometric and physiological data for women and men are presented. There were differences between the sexes for most factors except for age, trunk muscle endurance, and muscle fibre distribution.

VO_2_ increased with both speed and load in both women and men (Table 2). On all loads men have higher absolute VO_2_ values but lower values in percent of VO_2_max. Mean values for the 50 kg load at 5 km h^-1^ was 57 and 44 % of VO_2_max in women and men, respectively. There was no difference between the sexes in ELI20, but regarding ELI35 and ELI50 women had significantly higher values.

**Table 2.** VO_2_ in l min^-1^ (mean ± SD) and in percent of VO_2_max (mean ± SD) and ELI at walking speed 3 and 5 km h^-1^ carrying 20, 35 and 50 kg in women and men, respectively. *denotes p<0,05 ELI difference between women and men.

|  | *Women* |  |  | *Men* |  |  | *ELI*<br>*-differences* |
| --- | --- | --- | --- | --- | --- | --- | --- |
| *Speed/load* | *$\text{l min}^{-1}$* | *% $\text{VO}_{2\text{max}}$* | *ELI* | *$\text{l min}^{-1}$* | *% $\text{VO}_{2\text{max}}$* | *ELI* |  |
| **3 $\text{km h}^{-1}$** |  |  |  |  |  |  |  |
| 20 kg | $0.80 \pm 0.08$ | $23 \pm 2$ | $1.14 \pm 0.11$ | $1.02 \pm 0.15$ | $22 \pm 3$ | $1.13 \pm 0.13$ | 0.68 |
| 35 kg | $1.05 \pm 0.08$ | $26 \pm 3$ | $1.42 \pm 0.17$ | $1.19 \pm 0.12$ | $26 \pm 3$ | $1.32 \pm 0.14$ | 0.050* |
| 50 kg | $1.28 \pm 0.07$ | $39 \pm 4$ | $1.73 \pm 0.17$ | $1.42 \pm 0.16$ | $31 \pm 5$ | $1.57 \pm 0.21$ | 0.020* |
| **5 $\text{km h}^{-1}$** |  |  |  |  |  |  |  |
| 20 kg | $1.17 \pm 0.11$ | $32 \pm 3$ | $1.11 \pm 0.94$ | $1.37 \pm 0.18$ | $30 \pm 4$ | $1.15 \pm 0.97$ | 0.19 |
| 35 kg | $1.48 \pm 0.14$ | $45 \pm 4$ | $1.49 \pm 0.19$ | $1.63 \pm 0.18$ | $36 \pm 4$ | $1.33 \pm 0.14$ | 0.035* |
| 50 kg | $1.86 \pm 0,11$ | $57 \pm 5$ | $1.82 \pm 0.18$ | $1.97 \pm 0.16$ | $44 \pm 6$ | $1.60 \pm 0.16$ | 0.001* |

HR was significantly lower on all loads and both speeds in men compared to women. Carrying 50 kg at 5 km h^-1^ the mean HR value for women was 149 ±17 bpm and 120 ±19 bpm for men.

HLa was low on all loads. At 50 kg walking 5 km h^-1^ the mean HLa value was 2.00 ± 0.19 mM in women and 1.31 ± 0.57 mM in men (p<0.05).

RPE for general, back, leg and breathing fatigue was higher on all loads in women compared to men except for the lowest loads (20 kg, both speeds). RPE for back was highest in both men and women. At 50 kg and 5 km h^-1^ the RPE for back was 14.1 and 15.1 for men and women, respectively (p<0,05).

### ELI analyzes

#### ELI20

For ELI20, significant correlations were found between body mass (rho = 0.40) and BMI (rho = −0.44), but not for any other of the investigated factors. No significant difference between sexes was found in ELI20, and hence no meaningful mediation analysis could be performed.

#### ELI35

A significant difference between sexes for ELI35 was found (1.33 and 1.45, respectively, p<0.05). For ELI35 significant correlations were found for height (rho = −0.50), VO_2_max in l min^-1^ (rho = −0.37), leg strength (rho = −0.33) and for type IIa muscle fibre distribution (rho = 0.47). All candidates were tested for mediation, with a significant direct c-path between ELI35 and sex of −0.12 (p = 0.03). None of the candidates yielded an independent, significant mediation effect (all p>0.05) – see Table 3.

**Table 3.** Bootstrapped mediation (ab-) paths coefficients (95 % CI) for proposed mediators between sex and ELI35. Relation between sex and ELI35 after mediation is expressed as c´-path and corresponding p-value.

|  | *ab-coefficients* | *95% CI* | *c'-path* |
| --- | --- | --- | --- |
| Height (cm) | -0.124 | -0.27 to -0.02 | 0.00 (ns) |
| Body mass (kg) | -0.073 | -0.210 to 0.049 | -0.049 (ns) |
| BMI ($\text{kg m}^{-2}$) | 0.002 | -0.065 to 0.072 | 0.119 (ns) |
| $\text{VO}_2$ ($\text{l min}^{-1}$) | -0.064 | -0.244 to 0.076 | -0.058 (ns) |
| $\text{VO}_2$ ($\text{ml min}^{-1} \text{kg}^{-1}$) | 0.022 | -0.054 to 0.103 | -0.144 ($p < 0.01$) |
| Leg Strength (watt) | -0.053 | -0.200 to 0.057 | -0.068 (ns) |
| Trunk endurance (s) | 0.002 | -0.013 to 0.031 | -0.124 ($p < 0.03$) |
| Type I (%) | 0.014 | -0.007 to 0.082 | -0.140 ($p < 0.02$) |
| Type IIa (%) | -0.029 | -0.089 to 0.001 | -0.097 (ns) |
| Type IIx (%) | -0.014 | -0.073 to 0.042 | -0.111 (ns) |

#### ELI50

ELI50 was found to be lower in men, compared to women (1.61 and 1.82, respectively, p<0.05). Significant (p<0.05) bivariate correlations to ELI50 were found for height (rho =−0.63), body mass (rho = −0.77), BMI (rho = −0.58), VO_2_max in l min^-1^ (rho = −0.66) and leg muscle strength (rho = −0.67). No significant correlation were found for muscle fibre distribution, VO_2_max in ml kg_-1_ min_-1_, or trunk muscle endurance.

All candidates were tested for mediation between sex and ELI50 (Table 4). The direct effect of sex on ELI50 (c-path) was found to be 0.209 (p<0,05). Generally, body size, leg strength and VO_2_max in l min^-1^ all acted as mediators in uncontrolled analyses. Height, body mass and BMI all correlated strongly, and body mass provided the strongest mediation effect. When entering VO_2_ (l min^-1^), leg strength and body mass in a combined analysis (multimediator analyses), only body mass yielded a significant mediation effect (0.227, 95 % CI: 0.070 to 0.432) and a non-significant c´-path.

**Table 4.** Bootstrapped mediation (ab-) paths coefficients (95 % CI) for proposed mediators between sex and ELI50. Relation between sex and ELI50 after mediation is expressed as c´-path and corresponding p-value.

|  | *ab-coefficients* | *95% CI* | *c'-path* |
| --- | --- | --- | --- |
| Height (cm) | 0.154 | 0.023 to 0.320 | 0.063 (ns) |
| Body mass (kg) | 0.225 | 0.126 to 0.369 | -0.016 (ns) |
| BMI ($\text{kg m}^2$) | 0.071 | 0.015 to 0.118 | 0.140 ($<0.05$) |
| $\text{VO}_2$ ($\text{l min}^{-1}$) | 0.224 | 0.093 to 0.392 | -0.009 (ns) |
| $\text{VO}_2$ ($\text{ml min}^{-1} \text{kg}^{-1}$) | - 0.042 | -0.154 to 0.404 | 0.252 ($p < 0.01$) |
| Leg strength (watt) | 0.228 | 0.092 to 0.396 | -0.020 (ns) |
| Trunk endurance (s) | 0.002 | -0.035 to 0.011 | 0.209 ($p < 0.01$) |
| Type I (%) | -0.023 | -0.095 to 0.009 | 0.224 ($p < 0.01$) |
| Type IIa (%) | 0.009 | -0.011 to 0.082 | 0.195 ($p < 0.01$) |
| Type IIx (%) | -0.028 | -0.101 to 0.009 | 0.230 ($p < 0.01$) |

### Experience of carrying loads

The ELI between the 16 experienced and the 20 unexperienced participants were respectively: ELI20 1.08 ± 0.08 and 1.17 ± 0.09 (p<0.05); ELI35 1.32 ± 0.12 and 1.43 ± 0.19 (p = 0.058); and ELI50 1.60 ± 0.16 and 1.79 ± 0.07 (p<0.05).

### Accelerometry

There was no relation between neither body length and mass, carrying experience or sexes between number of counts neither in the x-, y- and z- direction nor in total vector counts on any load and speed.

## Discussion

The overall results of this study verify results for previous investigations (*e.g.* Goldman and Lampietro 1962, Pandolf et al. 1976, Soule et al. 1978, Christie and Scott, 2005). Energy expenditure increases with both speed and load. Oxygen uptake on all loads are higher in men than women due to average heavier body mass in men. Energy expenditure in relation to body mass (relative VO_2_) does not differ between the sexes on the different loads (data not shown). Åstrand (1967) has suggested that the highest load for a healthy all day physical work is about 40 % of VO_2_max. In this study this limit was nearly reached (39 %) by women for 50 kg load at the slower speed, 3 km h^-1^. At the higher speed, 5 km h^-1^, this limit was passed by women carrying 35 and 50 kg and by men when carrying 50 kg. It should be mentioned that when men carry 35 kg at 5 km h^-1^ the SD indicates that about a third of the men are stressed to 40 % of VO_2_max or more. Data on RPE values support this. These data on the energy expenditure during the different walking speeds and loads are in essence relatively similar to those earlier reported by Christie and Scott (2005). They used the same Åstrand health limit for prolonged work. The limit suggested by Åstrand deals mainly with all day manual industrial workloads and speeds and workloads higher than that may cause premature fatigue. Roy et al. (2012 and 2013) have shown that load and not the carrying time is the most important factor for overuse injuries. It should be mentioned that the HLa values in this study was low. This might partly be due to the short exercise time on each load, but other unknown factors cannot be excluded.

The ELI has been used as a proxy for load carrying capacity. For lighter loads (20 kg) there were no differences between the sexes, neither at 3 nor 5 km h^-1^ speed. Body mass and BMI were correlated to ELI20 at 5 km h^-1^ speed while there were no correlations to other factors studied. For the higher loads (35 and 50 kg) ELI was higher and differences between men and women were larger. Factors well known to be positively related to load carrying capacity such as body mass, body height and VO_2_max are confirmed in this study. On the other hand the non-related factors to load carrying capacity such as trunk muscle endurance and relative VO_2_max are novel and previously not described. With regard to muscle fibre distribution there is a mixed picture. At 35 kg and 5 km h^-1^ type IIa correlates to ELI but for ELI50 there seem to be no importance of muscle fibre distribution. Nor was muscle fibre composition a significant mediator for gender differences.

There were differences between the sexes in ELI values for both 35 and 50 kg loads but were these differences primarily due to the sex? Mediator analyses using ELI50 as an outcome and sex as the causal variable body size, leg strength and VO_2_max in l min^-1^ all acted as mediators in uncontrolled analyses. Height, body mass and BMI all correlated strongly but body mass provided the strongest mediation effect. When entering VO_2_max in l min^-1^, leg strength and body mass in a combined analysis, only body mass yielded a significant mediation effect. Thus ELI50 differed between sex mediated via differences in body mass.

An interesting questions is to what extend long term experience and, thus, carrying training, is important for carrying ability in relation to other studied factors with regard to the known extreme carrying performance among the short Sherpas and Tibetan Highlanders. ELI values of 20 kg and 50 kg at 5 km h^-1^ were significantly lower in the group of men and women with prolonged experience in carrying heavy loads compared to those without such an experience (with borderline difference for ELI35). None of the studied factors in this study could explain or mediated this difference. This result is supported by the finding in the study of Minetti et al (2006). The higher mechanical efficiency was also observed in the study by Bastien et al. (2005) and there in part explained by a lesser trunk oscillation both during downhill and uphill walking. The improved carrying capacity in our participants could be due to the same mechanism, since no other factor explained the ELI differences between carrying experienced and control participants. On the other hand, the measurement of accelerometry counts did not show any relation to any of the studied factors in this study. This might indicate that Sherpas extreme carrying capacity is, at least to some extent, due to a long time training experience, even from young ages, which far exceeds the carrying training experience in our group of carrying trained participants.

### Strength and limitation

Participants in present study were both trained and relatively untrained for carrying heavy loads. This limits the risk of positive selection. A strength is that several important factors were studied. The statistical approach with mediation analyses is also a strength in this study. All physiological data was collected with validated methods in a controlled environment. Walking on a soft carpet with some elevations mimicking track walking is as far as we know uniquely done in this study.

A limitation could be the method used for trunk endurance, but we have no indication that another method for evaluation of trunk endurance would correlate better with carrying ability.

### Statistical consideration

An alternative statistical approach to evaluate and analyse the carrying capacity between the sexes could have been to carry out a backward exclusion linear regression including sex, using VO_2_max (l min^-1^), leg strength and body mass as independent variables and ELI50 as dependent. Relative VO_2_max was not included, since body mass is already taken in consideration in the analyses. We performed such analysis. The result showed that body mass was the only variable left in the final model (data not shown). This indicate that the results are not dependent on the statistical model used.

### Perspectives and conclusion

The findings in this study have practical consequences. First of all sex is not an important factor, even if mediator analyses showed a small c´ path after adjusting for other relevant factors. For selection of participants for work or tasks involving carrying heavy loads, in this study represented by a load of totally 35 and 50 kg, the most important factor to consider is body mass. The body weight and height have evidently some importance, but they are to a large part not changeable. Factors that can be changed by physical training such as muscle strength and trunk endurance seem to be of lesser importance. In addition neither relative nor absolute VO_2_max nor muscle fibre distribution seem to predict load carrying performance. But the analyses of carrying experience show that the importance of body mass can be abolished by long term carrying training.

